# Genomic signatures of selection in *Anopheles funestus* reveal shared and population-specific adaptive variation across African populations

**DOI:** 10.64898/2026.05.07.723524

**Authors:** Helga Modukpe Saizonou, Abdoulaye Sadio, Harouna Dit Massire Soumare, Kelly L. Bennett, Majidah Adiamoh, Eniyou Oriero, Jon Brenas, Anastasia H. Koutoucheva, Mamadou O. Ndiath, Alistair Miles, Annette Erhart, Umberto D’Alessandro, Chris S. Clarkson, Benoit S. Assogba, Janet Midega, Christophe Antonio Nkondjio, El Hadji Amadou Niang, Eugène Kaman Lama, Olaitan Omitola, Alfred Amambua-Ngwa

## Abstract

Insecticide resistance in *Anopheles funestus* threatens malaria vector control across sub-Saharan Africa, yet genomic signatures of selection across geographically structured populations remain poorly understood. We analysed whole-genome sequence data from 635 *An. funestus* mosquitoes from West (Senegal, Guinea, Nigeria), Central (Cameroon), and East Africa (Kenya) to characterise population structure and identify targets of recent positive selection.

Population genomic analyses revealed strong differentiation between East African and West/Central African populations, with finer-scale structuring within West Africa. Genome-wide selection scans using H12 identified sweep regions on chromosome arms 2RL and 3RL, overlapping a cytochrome P450 cluster and the gamma-aminobutyric acid (GABA) receptor locus respectively. iSAFE prioritised candidate variants within these sweeps: non-synonymous substitutions in CYP6A14 were identified at high frequencies in Guinea, Nigeria, and Cameroon, while population-restricted variants implicated octopamine receptor genes on 2RL and the GABA receptor on 3RL. Diplotype clustering and copy number variation analyses confirmed the causal role of candidate variants, embedded within extended haplotypes of reduced heterozygosity consistent with recent positive selection.

These findings demonstrate that adaptive evolution in *An. funestus* reflects both shared and population-specific selective processes shaped by geography and ecological context. Whereas selection on detoxification pathways appears widespread, localised signals in neuromodulatory loci, including the GABA receptor and octopamine-related genes, reveal that biological systems beyond metabolic resistance contribute to mosquito adaptation. The convergence of selective signals across these gene classes highlights neuromodulatory pathways as potential complementary targets for next-generation vector control strategies.

## Introduction

Malaria remains one of the most important public health challenges worldwide, with most cases and deaths occurring in sub-Saharan Africa (WHO, 2025). The disease is transmitted by some female *Anopheles* mosquitoes, making vector control an important factor in malaria prevention strategies (WHO, 2025). Over the past two decades, the large-scale deployment of insecticide-based interventions such as long-lasting insecticide-treated nets (LLINs) and indoor residual spraying (IRS) has significantly reduced malaria transmission in many endemic regions (WHO, 2025). However, the emergence and rapid spread of insecticide resistance in mosquito populations threaten the effectiveness of these interventions.

Insecticide resistance has been extensively studied in several malaria vectors, particularly members of the *Anopheles gambiae* complex, where decades of research have identified multiple resistance mechanisms including target-site mutations, metabolic detoxification, and behavioural adaptations (Hancock et al., 2024; Riveron et al., 2018). In contrast, although *Anopheles funestus* is one of the major malaria vectors across Africa and can sustain malaria transmission even in areas when other vectors decline during the rainy season (Adja et al., 2011; Akoton et al., 2023; Djamouko-Djonkam et al., 2020), the evolutionary mechanisms underlying its adaptation to insecticide pressure remain relatively less understood.

*An. funestus* is typically associated with permanent or semi-permanent water bodies and is widely distributed across diverse ecological settings, including humid forest regions and savannah environments. These ecological differences can influence mosquito population dynamics, exposure to vector control interventions, and local selection pressures (Dia et al., 2013; Nambunga et al., 2020). Previous studies have shown that the resistance phenomenon in this specie is largely driven by metabolic mechanisms involving the overexpression of detoxification enzymes such as cytochrome P450 monooxygenases, esterases, and glutathione-S-transferases (Atoyebi et al., 2020; Sangba et al., 2016; Suh et al., 2023). Although most work has focused on metabolic detoxification, other physiological systems, including neuromodulatory pathways, could contribute to adaptation to insecticide pressure and environmental stress in this species. Neuromodulators such as octopamine and GABA regulate key processes including neuronal excitability, locomotion, feeding behaviour, and stress responses, all of which can influence how mosquitoes respond to insecticide exposure (Ellis, Alampounti, et al., 2025; Farooqui, 2012; Freeman et al., 2024). For instance, modulation of neural activity may affect behavioural avoidance, recovery following sublethal exposure, or sensitivity to neurotoxic compounds. While these mechanisms remain less well characterized in *An. funestus*, they provide a plausible link between neural regulation and resistance phenotypes. As such, multiple biological systems may contribute to adaptation in *An. funestus*. However, the broader genomic signatures associated with these processes, and how they are shaped by selective pressures across different African populations, remain poorly characterised. From an evolutionary perspective, strong selective pressures such as widespread insecticide use can leave detectable footprints in mosquito genomes (Calla et al., 2021; Weedall et al., 2020a). Beneficial mutations that enhance survival under insecticide exposure can rapidly increase in frequency, generating genomic signatures known as selective sweeps. Identifying such signatures provides valuable insights into the evolutionary dynamics shaping vector adaptation and can reveal candidate genes and pathways involved in resistance. Population genomic approaches have increasingly been used to detect these adaptive signals in malaria vectors, helping to uncover the genetic basis of insecticide resistance and environmental adaptation (Boddé et al., 2025; Hancock et al., 2024). While these approaches have provided important insights, how sweeps are distributed across geographically and ecologically diverse *An. funestus* populations, and the extent to which adaptive variants are shared or locally restricted, remains poorly understood. Understanding these patterns is critical, as populations experiencing similar ecological conditions may evolve comparable adaptive responses despite large geographic distances, while neighbouring populations may follow distinct evolutionary trajectories under different local pressures.

In this study, we investigated the population genomic structure and signatures of selection in *An. funestus* using publicly available whole-genome datasets from multiple African regions. By analysing populations from West Africa (Senegal, Guinea, and Nigeria), Central Africa (Cameroon), and East Africa (Kenya), we aimed to identify genomic regions under positive selection and prioritise candidate variants driving adaptive evolution. Integrating genome-wide selection scans with population structure analyses and variant prioritisation, we uncover both shared and population-specific selective signals implicating detoxification enzymes and neuromodulatory pathways, highlighting the complexity of adaptation in this species and revealing potential targets beyond classical insecticide resistance mechanisms for future vector control strategies.

## Materials and Methods

### Mosquito collection

This study analysed *An. funestus* data generated by the MalariaGEN Vector Observatory. The *An. funestus* Mosquitoes were collected in Guinea, Nigeria, and Senegal from West Africa, and Cameroon from Central Africa, while additional open-source samples, from Kenya in East Africa were included; yielding a total of 635 samples (**Supplementary file 1**).

In Senegal, mosquitoes were collected between 2020 and 2022 from Kaolack, a region in the central western part of the country; Kédougou and Kolda in the southern part near the borders with Guinea-Bissau and Mali; Louga; and Saint-Louis in the northern part of the country towards Mauritania. In Guinea, collections were carried out in 2022 in Kankan and Nzérékoré. Kankan is in the northeastern part of the country and serves as a major commercial and transport hub in Upper Guinea, where the Niger River originates. Nzérékoré, by contrast, is located in the forested highlands of southern Guinea bordering Sierra Leone and Liberia. In Nigeria, mosquitoes were collected in 2018 from Ogun and Oyo states, two neighbouring states in the southwestern region of the country, an area characterized by intensive agricultural activities and influenced by the Ogun River. In Cameroon, collections were conducted in the southern part of the country in 2020 from the Southwest equatorial forest region. In Kenya, mosquitoes were collected earlier, between 2010 and 2012, in Kilifi County, located in the southeastern part of the country along the Indian Ocean coast and considered as an endemic zone (Polo et al., 2025) (**Figure 1-A**).**Sequencing and sequence data processing**

**Figure 1.**
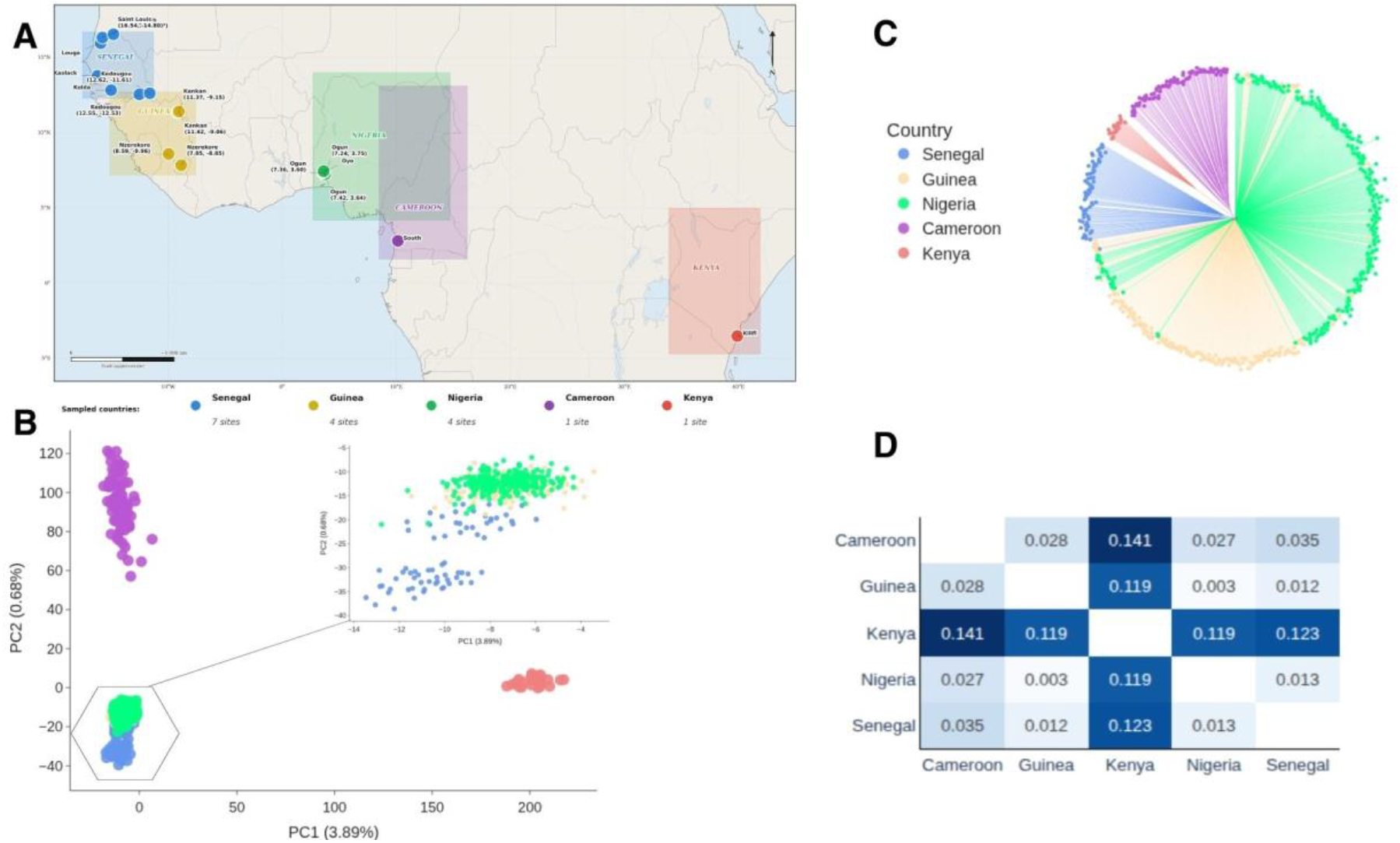
Sampling locations and population structure of An. funestus across Africa. (**A**) Geographic distribution of sampled populations in West, Central, and East Africa. Each country is colour coded as indicated in the legend, and dots represent individual collection sites. (**B**) principal component analysis (PCA) on variants from the inversion-free region of chromosome arm 2RL (57,604,655–90,000,000). The plot shows the first two principal components, with samples coloured by country of origin. (**C**) Unrooted neighbor-joining tree (NJT) constructed from the same region with samples coloured according to country of origin. (**D**) Pairwise genetic differentiation (FST) among populations calculated from the same genomic region and visualised as a heatmap. Values represent pairwise FST estimates between countries.

Sequencing and primary data processing were conducted following previously published protocols described in (Boddé et al., 2025) and as implemented in the MalariaGEN pipelines available from the MalariaGEN GitHub repository (https://github.com/malariagen/pipelines/). Detailed descriptions of the sequencing, alignment, and variant calling procedures are as previously described (Boddé et al., 2025). Libraries were prepared using a standard workflow for the NEBNext Ultra II DNA Library Prep Kit for Illumina and sequenced on the Illumina HiSeq X10 platform using paired-end 150 bp reads, targeting approximately 30× coverage per individual.

Read alignment and genotyping followed the MalariaGEN alignment and genotyping pipelines with default parameters (https://github.com/malariagen/pipelines/). Sequencing reads were aligned to the AfunGA1 reference genome using *bwa mem*, and alignments were post-processed using standard tools to merge lanes, mark PCR duplicates, and realign reads around indels. Variant calling was performed using GATK UnifiedGenotyper, considering all possible substitutions at non-ambiguous reference genome sites. Variant call format (VCF) files were converted to Zarr format using *scikit-allel* for downstream analyses.

Quality control (QC) and site filtering were performed following the criteria defined in (Boddé et al., 2025) and as described in the MalariaGEN Vector Data user guide (https://malariagen.github.io/vector-data/ag3/methods.html). These procedures evaluated sequencing coverage, genome completeness, divergence from the reference genome, contamination levels, sex concordance, and sample relatedness. Two site-level filters were applied: a static-cutoff (sc) filter for most analyses and a decision-tree (dt) filter for haplotype-based analyses, consistent with the filtering strategy used in those studies.

### Population structure and patterns of genetic diversity

Population structure of the Guinea and Nigeria population, together with Senegal in West Africa, Cameroon in Central Africa, and Kenya in East Africa, was investigated using principal component analysis (PCA), an unrooted neighbour-joining tree (NJT), and pairwise genetic differentiation (FST) across the 2RL: 57,604,655–90,000,000 region. This region was chosen because it is not affected by large structural rearrangements and is therefore inversion-free. PCA and NJT analyses were conducted using the *malariagen_data* Python API (v15.5.0) (https://malariagen.github.io/malariagen-data-python/latest/Af1.html), with 100,000 biallelic SNPs equally distributed across the region. PCA enabled visualization of the overall genetic structure and the distribution of samples, while the NJT helped distinguish how samples from each location clustered relative to one another. To further quantify genetic differentiation among populations, pairwise FST values were computed using the *af1*.*pairwise_average_fst* function from the *malariagen_data* Python API for all population pairs within the same genomic region, allowing direct comparison of differentiation levels among countries. These analyses provided a complementary quantitative assessment of population structure alongside PCA and NJT.

In addition to multivariate analyses, population genetic summary statistics were computed to quantify patterns of genetic diversity. Nucleotide diversity (π) and Tajima’s D were estimated using the same inversion-free 2RL region (57,604,655–90,000,000), ensuring direct comparison between inferred population structure and levels of genetic variation. To quantify extended homozygous tracts within individuals, runs of homozygosity (ROH) were inferred independently across the entire 2RL chromosome arm using the hidden Markov model (HMM) implemented in scikit-allel. ROH segments shorter than 100 kb were excluded. For each sample, the number of ROH segments and the fraction of the 2RL arm contained within ROH (F_ROH) were calculated. ROH analyses were performed separately by country and summarized to evaluate differences in the extent and distribution of homozygous tracts across populations.

### Genome-wide selection scans and population differentiation

Genome-wide selection scans (GWSS) were conducted independently for each country’s population across all chromosome arms using the *h12_gwss* function implemented in the *malariagen_data* Python API. Prior to GWSS, country-specific H12 calibration curves were generated using the *plot_h12_calibration* function. Window sizes were selected such that the 95th percentile of H12 values was below 0.1, accounting for differences in sample size and haplotype diversity among countries. Based on these calibrations, GWSS statistics were computed and plotted, and genomic regions showing pronounced H12 peaks were identified for downstream analyses. To assess patterns of population differentiation within candidate regions, windowed FST was computed using the Weir and Cockerham estimator as implemented in *allel*.*windowed_weir_cockerham_fst* (scikit-allel.readthedocs.io/en/stable/stats/fst.html), using non-overlapping windows of 10 kb. This allowed visualization of local variation in genetic differentiation along genomic coordinates. To assess haplotype sharing between population pairs within each candidate region, pairwise H1X statistics were computed using the *h1x_gwss* function implemented in the *malariagen_data* Python API, with a window size of 2,000 bp and a minimum cohort size of 15 individuals. For each candidate region, H1X values were extracted within the defined genomic interval and summarized as the median across windows, then visualized as pairwise heatmaps. High H1X values between two populations indicate that the most frequent haplotype is shared across both populations, consistent with a common selective sweep.

### Haplotype-based SNP prioritization using iSAFE

Genomic regions showing elevated genome-wide selection statistics (GWSS) were further examined by extracting genotype data for all genes falling within each candidate region. Haplotypes data were retrieved using the *haplotypes* function from the *malariagen_data* Python API. Samples were stratified into cohorts defined by site of collection, country, and pooled multi-country groupings, and analyses were performed separately for each cohort. Candidate regions were restricted to a maximum length of 6 Mbp. For each region and cohort, diploid genotypes calls were subset by cohort and converted into haplotype matrices using the *to_haplotypes* function implemented in *scikit-allel*, yielding two haplotypes per individual. Variant genomic positions were extracted alongside the haplotypes and used to order SNPs by physical position along the chromosome. The resulting position-sorted haplotype matrices were used to generate region-specific haplotype input files for downstream iSAFE analyses.

To prioritize individual variants within these regions that could underlie the observed selective sweeps, we applied the iSAFE model (Akbari et al., 2018), a haplotype-based method that ranks SNPs based on patterns of haplotype structure and allele frequency consistent with recent positive selection. iSAFE analyses were restricted to genomic regions ≥200 kb in size with sufficient SNP density, excluding regions with ≤100 SNPs as flagged by the iSAFE implementation (https://github.com/alek0991/iSAFE/blob/master/isafe/isafe.py). iSAFE scores were computed for SNPs within each region, and variants with iSAFE scores ≥ 0.25 were retained for downstream analyses. These high-scoring variants were considered prioritized candidates and mapped to their corresponding genes using genomic coordinates from the reference genome annotation.

### Gene-level interpretation of selection signals

For each prioritised SNP, the predicted functional annotation (e.g., synonymous, non-synonymous, or intergenic) was recorded. Because iSAFE analyses were conducted separately for each country as well as across all populations combined, prioritised variants were subsequently classified according to their geographic distribution. SNPs detected in the combined analysis and consistently identified across multiple populations were considered shared candidate signals, whereas variants detected independently in more than one country were considered shared among subsets of populations.

To further characterize genetic structure within candidate regions, diplotype-clustering analyses were performed using genotypes data spanning genomic intervals containing prioritized SNPs. These analyses allowed visualization of patterns of genetic similarity among individuals within and across populations, while also capturing variation in heterozygosity across the region. This approach provides a more complete representation of individual genetic variation, including signals that may be difficult to resolve through phased haplotypes alone (Nagi et al., 2024). In parallel, these same genomic regions were examined for evidence of copy number variation (CNV) to assess whether structural variation co-occurred with SNP-based signatures of selection.

## Results

### Population structure revealed strong regional genetic differentiation in *Anopheles funestus*

PCA revealed clear genetic differentiation among *An. funestus* populations from West, Central, and East Africa, based on variants from the inversion-free region of chromosome arm 2RL (57,604,655–90,000,000) (**Figure 1-B**). The first two axes of variation separated Kenyan and Cameroonian samples from the West African populations. Within the latter, samples from Senegal, Guinea, and Nigeria showed partial as additional substructure was evident overlap within Senegal, where individuals from Kolda and some from Kédougou formed distinct clusters separated from other Senegalese sites and from Guinea and Nigeria (**Supplementary file 2**). The NJT constructed from the same genomic region showed a clustering pattern consistent with the PCA results (**Figure 1-C)**. Samples were grouped largely according to country and geographic proximity, with Kenyan and Cameroon mosquitoes forming different, distinct branches, while mosquitoes from West African populations clustered more closely together.

Pairwise FST estimates further supported these observed population structure patterns (Figure 1-D). Differentiation was highest between Kenyan samples and all other populations (0.12-0.14), while Cameroon also showed moderate differentiation from West African populations (0.02 - 0.035). In contrast, lower FST values were observed among West African populations, particularly between Guinea and Nigeria (0.003), indicating closer genetic similarity. Senegal displayed intermediate levels of differentiation, consistent with the substructure observed in the PCA and NJT.

### Patterns of genetic diversity and run of homozygosity revealed regional contrasts among *Anopheles funestus* populations

We computed genetic diversity across all populations using nucleotide diversity (π), Tajima’s D, and runs of homozygosity (ROH) (**Figure 2**). Nucleotide diversity (**Figure 2-A**) was broadly similar across West African populations, (π = 0.027 to 0.030) with one notable exception: samples from Louga population showed reduced nucleotide diversity, compared to Cameroon population. Kenya samples from Kilifi exhibited lower nucleotide diversity (~0.023), clearly differentiating them from all other populations analysed.

**Figure 2.**
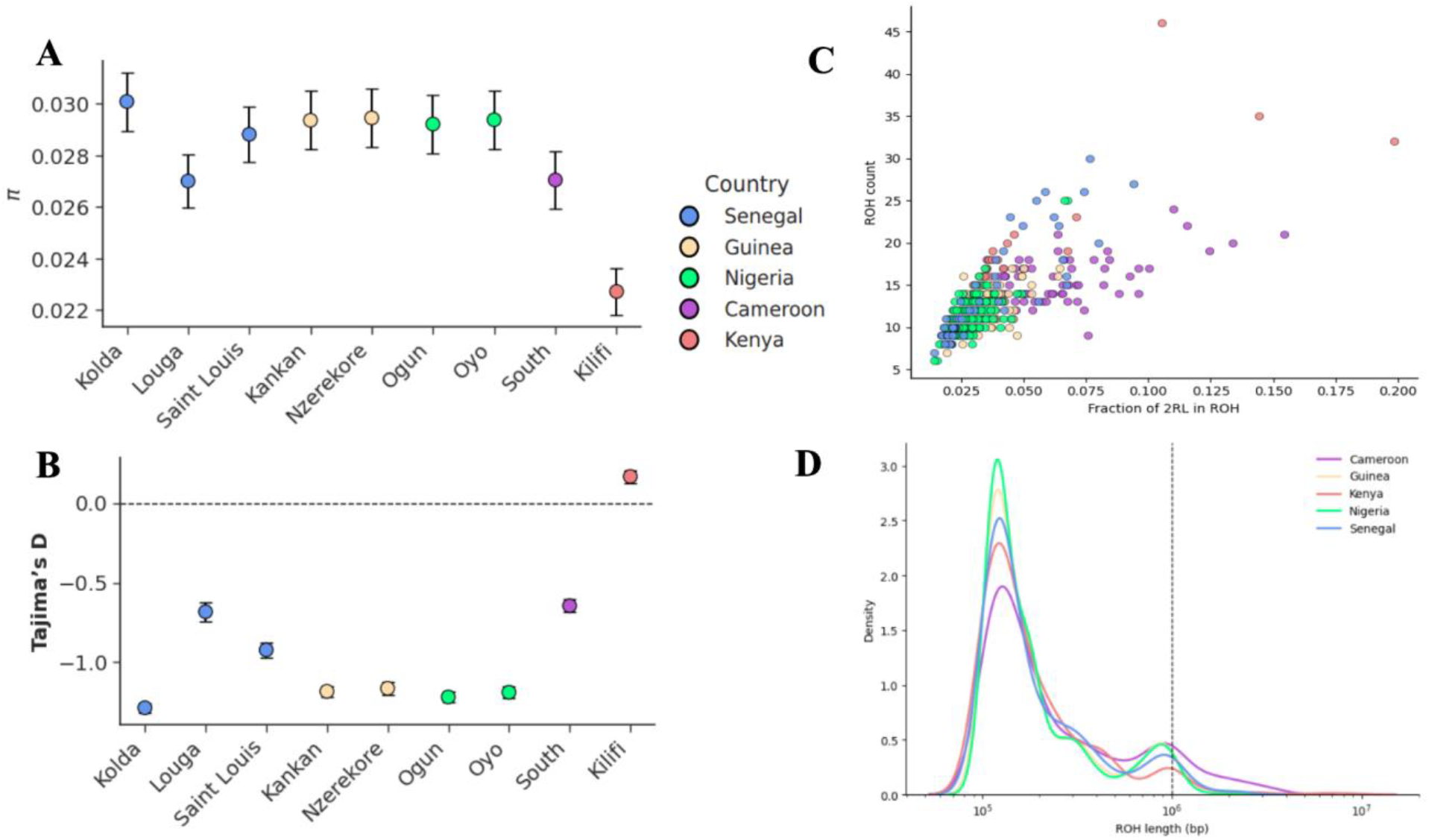
Genetic diversity and runs of homozygosity (ROH) across *An. funestus* populations. (A) Nucleotide diversity (π), and (B) Tajima’s D, are shown with point estimates and corresponding confidence intervals for each population, coloured by country, with the sampling locations shown along the x-axis. The dashed horizontal line in panel (B) marks neutrality (D = 0). (C) Relationship between the fraction of chromosome arm 2RL contained within ROH (F_ROH) and the number of ROH segments per individual. Each point represents a single individual and is color-coded by country. (D) Distribution of ROH segment lengths across populations on chromosome arm 2RL. Density curves show the distribution of ROH lengths (log scale). The dashed vertical line indicates 1 Mb, separating shorter from longer homozygous tracts.

Tajima’s D (**Figure 2-B**) values were predominantly negative across West and Central African populations, with values ranging from moderately (~-0.5) to strongly (~-1.5) negative depending on sampling location. Kenyan samples displayed a positive value of Tajima’s D (~0.5) contrasting sharply with all other populations.

ROH was inferred across the entire 2RL chromosome arm. Both the number of ROH segments per individual and the fraction of 2RL contained within ROH (F_ROH) were examined (**Figure 2-C**). Most individuals from Senegal, Guinea, and Nigeria clustered at lower F_ROH values (approximately 0.02–0.06), while Cameroon and Kenya displayed a broader distribution, with multiple individuals exceeding F_ROH = 0.10. Kenyan samples contained the highest F_ROH values and ROH counts observed. The distribution of ROH tract lengths (**Figure 2-D**) was dominated by short tracts (~10^5 bp) across all populations. The upper tail revealed population specific patterns: Cameroon and Kenya showed a greater density of ROH tracts exceeding 1 Mb (vertical dashed line), while West African populations were largely enriched for shorter tracts.

### Population-structured selective sweeps with limited haplotype sharing highlighted parallel adaptation across African populations

Genome-wide selection scans (GWSS) based on the H12 statistic were conducted for each country across chromosome arms 2RL, 3RL, and X, and then jointly visualised to enable direct comparison (**Figure 3**). Multiple regions on chromosome arm 2RL, showed high H12 values, including 2RL:8.0–9.85 Mb, 40–44 Mb, 76.0–76.84 Mb, and 80.48–80.83. Some signals were shared across countries, whereas others were restricted and differed in peak magnitude (**Supplementary file 3**). Genes content and annotations for these regions are provided (**Supplementary file *4***).

**Figure 3.**
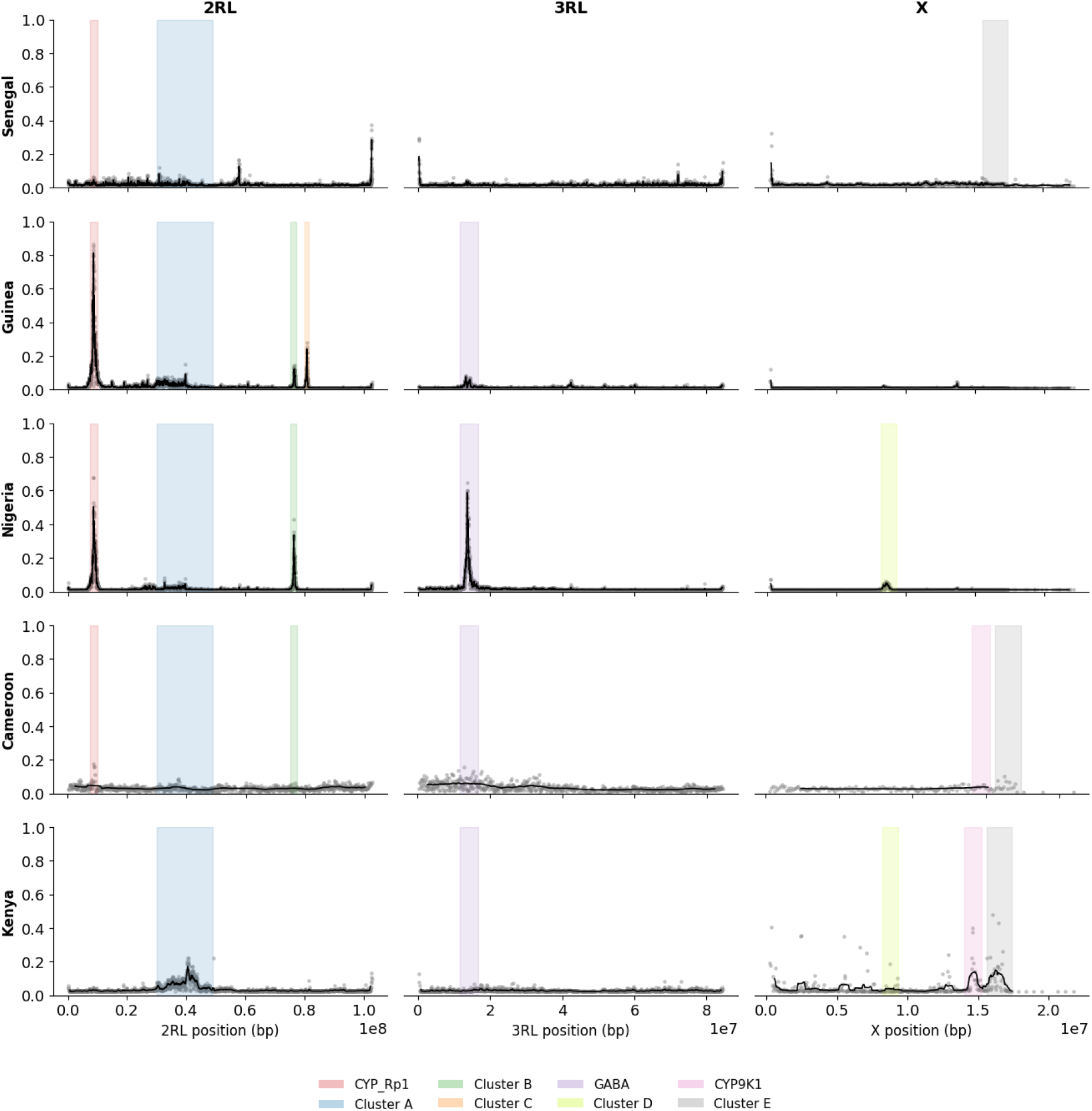
Genome-wide selection scans across chromosome arms 2RL, 3RL, and X Genome-wide selection scans based on the H12 statistic for *An. funestus* populations from Senegal, Guinea, Nigeria, Cameroon, and Kenya. Each row corresponds to a population, and points represent window-based H12 estimates along genomic coordinates. Shaded regions indicate candidate selective sweep regions identified from peak signals.

The strongest signal on 2RL mapped to the proximal region (8.0–9.85 Mb), which contains a cluster of cytochrome P450 genes along with other genes, including sodium/calcium exchanger 3 (LOC125764682), esterase E4-like (LOC125764700) (**Figure 3**). Local Fst profiles across this region showed elevated differentiation centred on the sweep peak, consistent with divergence in allele frequencies among populations (**Figure 4**). However, haplotype sharing measured using H1X remained low across most population pairs, with limited sharing between Guinea and Nigeria. This combination of strong sweep signal and low haplotype sharing supports repeated selection on similar genomic targets but involving distinct haplotypes, rather than the spread of a single adaptive variant.

**Figure 4.**
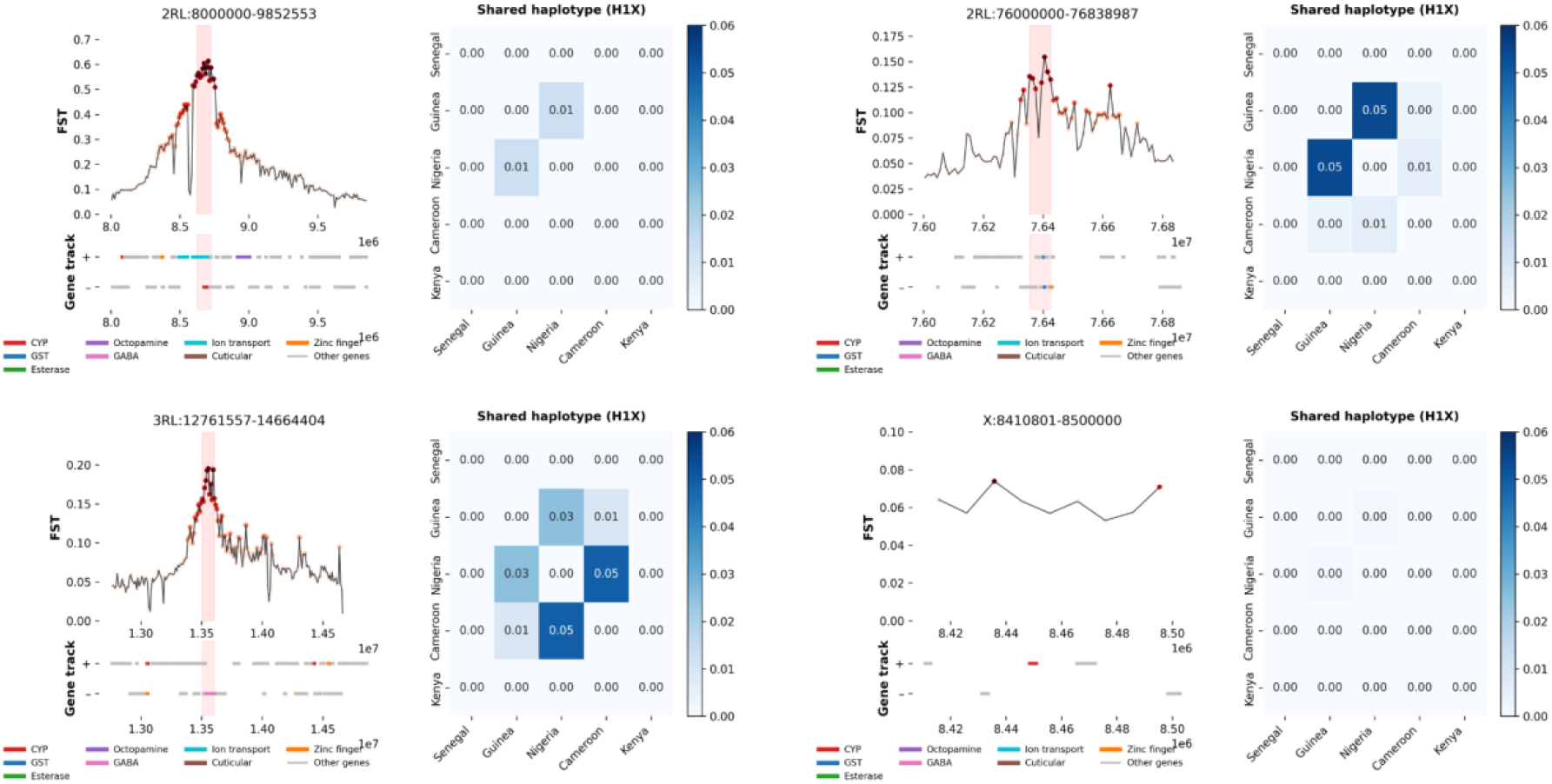
Patterns of genetic differentiation across candidate genomic regions. Genomic windows showing pairwise population differentiation (FST) and shared haplotype structure (H1X) across selected regions identified by genome-wide scans. For each region, the upper panel displays FST values along genomic coordinates, with the underlying gene annotation track indicating the position of candidate genes. The corresponding heatmap summarizes H1X values among populations (Senegal, Guinea, Nigeria, Cameroon, and Kenya), where higher values indicate greater sharing of the most frequent haplotype between population pairs, consistent with a common selective sweep.

The region at 2RL:40–44 Mb showed elevated H12 values with moderate differentiation across West and Central African populations, while Kenya remained distinct (**Figure 3**). H1X values across this region were uniformly low, indicating limited haplotype sharing and supporting a pattern of population-specific or weakly shared selective signals (**Supplementary file *5***).

At 2RL:57.63–58.0Mb, the high H12 values overlapped a GATA zinc finger gene, yet both FST and H1X values indicated minimal differentiation and negligible haplotype sharing across populations, suggesting that this signal may reflect weaker or more diffuse selective process (**Supplementary file *5***). Similarly, the region at 2RL:80,489–80.83, where H12 peaks were most pronounced in Guinea (**Supplementary file *5*)**, showed low haplotype sharing and limited differentiation, indicating that the signal is largely population-specific and not associated with a shared adaptive haplotype.

In contrast, the interval at 2RL:76.0–76.84Mb combined elevated H12 values with localised increases in differentiation, driven primarily by comparison involving Kenya (**Figure 3**). H1X values indicated limited sharing between East African and West/Central African populations, while modest sharing was observed among West African populations (**Figure 4**). Genes in this region included members of the glutathione S-transferase (GST) family, serine protease, CLIP-domain genes, and collagen-related genes, suggesting that both detoxification and immune-related pathways may contribute to adaptive variation in this interval.

On chromosome arm 3RL, a prominent H12 signal was detected at 12.76–14.66Mb (**Figure 3**). This region encompasses both cytochrome P450 genes and the γ-aminobutyric acid (GABA) gene. The GWSS signal was strongest in Nigeria and was also present in Guinea and Cameroon. Local FST profiles showed elevated differentiation centred on the GABA locus, and H1X analysis revealed moderate haplotype sharing among West and Central African populations (0.03-0.05), but no sharing with Kenya, indicating a partially shared sweep within West/Central Africa combined with regional divergence. (**Figure 4**).

On the X chromosome, GWSS identified fewer and generally weaker H12 peaks (**Figure 3**). A notable signal was observed at 8.41–8-5 Mb, overlapping the Cyp9k1 gene. Thedifferentiation in this interval was modest, and H1X values remained low across all population pairs, indicating minimal haplotype sharing between regions. This pattern suggests that adaptive signals on the X chromosome are either more recent, weaker, or more population-specific compared to those observed on autosomes. (**Figure 4**). Outside this interval, H12 values and FST estimates across the X chromosome remained low for most population comparisons.

### Prioritisation and characterisation of candidate variants within selective sweep regions

Haplotype-based SNP prioritisation using iSAFE identified candidate variants within the two strongest selective sweep regions detected by genome-wide H12 scans, located on chromosome arms 2RL (8.0–9.85 Mb) and 3RL (12.76–14.66 Mb). These regions correspond respectively to the cytochrome P450 gene cluster on 2RL and the GABA receptor region on 3RL (**Supplementary file *6***).

Within the 2RL sweep region (2RL:8,000,000–9,852,553), several high-scoring iSAFE variants (iSAFE score > 0.7) were detected across all populations (**Supplementary file *6***). These variants mapped to the sodium/calcium exchanger 3 (LOC125764682), esterase E4-like (LOC125764700), and cytochrome P450 6a14 (LOC125764705) genes. Variants identified in the sodium/calcium exchanger 3 and esterase E4-like genes were located within transcript regions and were predicted to have modifier effects. In contrast, several variants within cytochrome P450 6a14 corresponded to non-synonymous substitutions (L261S, F253I and I94L). The L261S mutation was fixed in Nigeria, Cameroon, and Kenya and nearly fixed in Guinea (~0.97), while occurring at lower frequency in Senegal (~0.44). Similarly, F253I was observed at near fixation in Guinea, Nigeria, and Cameroon (~0.95–0.99), but occurred at lower frequencies in Kenya (~0.18) and Senegal (~0.23). The I94L mutation was fixed in Nigeria and near fixation in Guinea, Cameroon, and Kenya, while remaining less frequent in Senegal (~0.28). Diplotype clustering across the region 2RL:8,915,154–9,064,309 (***Supplementary file 8***) that encompassed those genes, revealed distinct groupings characterized by extended tracts of low heterozygosity, consistent with signatures of recent positive selection. Notably, regions of reduced heterozygosity overlapped with genomic segments where cytochrome P450 6a14 variants (I94L, F253I, L261S) were fixed or near fixation, indicating that these mutations are embedded within dominant diplotype backgrounds. In addition, copy number variation (CNV) affecting several cytochrome P450 genes, including LOC125764712 (cytochrome P450 6a8-like), LOC125764714 (probable cytochrome P450 6a14), and LOC125764726 (probable cytochrome P450 6a13) was detected in populations exhibiting low heterozygosity (***Supplementary file 8*, Table 1**).

**Table 1.** Copy number variation (CNV) frequencies within the cytochrome P450 gene cluster on chromosome arm 2RL.

| Gene | Cameroon | Guinea | Kenya | Nigeria | Senegal |
| --- | --- | --- | --- | --- | --- |
| probable cytochrome P450 6a14 (amp) | 0 | 0.99 | 0 | 0.28 | 0 |
| cytochrome P450 6a8-like (amp) | 0 | 0.96 | 0 | 0.25 | 0 |
| probable cytochrome P450 6a13 (amp) | 0.96 | 0.06 | 0.79 | 0.39 | 0.71 |
| Cytochrome P450 (del) | 0 | 0.89 | 0 | 0.06 | 0 |
| esterase E4-like (del) | 0.26 | 0.88 | 0.07 | 0.11 | 0.36 |
| Cytochrome P450 (amp) | 0 | 0.02 | 0 | 0.2 | 0.68 |
| esterase E4-like (amp) | 0.59 | 0.02 | 0.18 | 0.03 | 0.2 |
| cytochrome P450 6a22-like (amp) | 0.06 | 0.24 | 0.11 | 0.31 | 0.23 |
| cytochrome P450 6a8-like (del) | 0.03 | 0 | 0.18 | 0.02 | 0.14 |
| probable cytochrome P450 6a13 (del) | 0 | 0.02 | 0 | 0 | 0 |

Within the same region (2RL:8,000,000–9,852,553), country-specific variants identified by iSAFE (Supplementary file 9) were also detected in additional genes. In Guinea, Cameroon, and Nigeria, prioritized variants were observed in the uncharacterized gene LOC125764675, while in Nigeria, additional variants were identified within the octopamine receptor beta-2R gene. Diplotype clustering across the interval 2RL:8,915,154–9,064,309 (**Supplementary file *9***), encompassing these two loci, revealed subsets of individuals in Guinea and Nigeria exhibiting extended tracts of low heterozygosity, consistent with localized signatures of positive selection. Notably, no copy number variation (CNV) was detected within these genes. However, a population-specific non-synonymous mutation (G111R) in LOC125764708_t3 (octopamine receptor beta-2R) was identified exclusively in Nigerian samples, coinciding with the low-heterozygosity cluster, while being absent in the Guinea samples with the same characteristics (***Supplementary file 8***).

Within the 3RL sweep region (3RL:12,761,557–14,664,404), prioritized variants were detected primarily within the gamma-aminobutyric acid (GABA) receptor gene and a probable nucleoporin gene (Nup54) (**Supplementary file *6*, Supplementary file *7***). These variants were predominantly detected in Nigeria and Cameroon, whereas weaker or no signals were observed in the other populations. All identified variants were located within transcript regions of the corresponding genes. No additional strong population-specific iSAFE signals were detected across other populations within this region, with the exception of a limited number of variants identified in Cameroon. Diplotype clustering across the region 3RL:13516317-13612792 encompassing these genes (**Supplementary file *9***), revealed a subset of individuals from Nigeria characterized by extended tracts of low heterozygosity. Within this group, a non-synonymous mutation (T345S) in the GABA receptor gene, with an iSAFE score of 0.25 across populations, was present in individuals exhibiting reduced heterozygosity, suggesting that it may be associated with the selective signal observed in this population. Further, no evidence of CNV was detected in these genes.

## Discussion

This study set out to characterise the genomic basis of adaptation in *An. funestus* across geographically and ecologically diverse African populations, and the results reveal a complex picture of both shared and population-specific evolutionary responses. By integrating genome-wide variation across West, Central, and East Africa, we provide new insight into the evolutionary processes shaping patterns of genetic differentiation and adaptive variation in this major but understudied malaria vector.

Our primary analyses focused on two recent datasets from collections of vectors from Guinea and Nigeria, both from West Africa, but separated by ~2000KM. Additional population included Senegal, neighbouring Guinea to the North and Cameroon neighbouring Nigeria to the East. As these population were contiguous between West and Central Africa, we introduced available Kenyan data for contrast. Population structure analyses affirmed the genetic distance of the Kenyan population from Kilifi and the Cameroonian population from the south, both of which formed distinct genetic clusters compared to West African populations. This pattern was consistently supported by PCA and NJT analyses, indicating substantial genetic differentiation across these regions. Similar structuring has been reported in *An. funestus*, where populations from East and Central Africa are genetically distinct from those in West Africa, suggesting limited gene flow at the continental scale. Such differentiation is often attributed to major geographic and ecological barriers, including the Congo Basin forest and the Great Rift Valley, which can restrict mosquito dispersal and promote regional divergence (Djuicy et al., 2020; Koekemoer et al., 2006). In Central Africa, forested environments have been shown to structure mosquito populations and reduce connectivity with surrounding regions (Michel et al., 2006). In addition, patterns of population structure in Cameroon have been linked to isolation-by-distance effects coupled with the non-random distribution of chromosomal inversions along environmental gradients, where shifts in inversion frequencies across ecological clines can generate hidden substructure and contribute to genetic differentiation even across relatively short distances (Michel et al., 2006). Moreover, the Great Rift Valley has also been proposed as a major geographic barrier limiting gene flow between eastern and western populations. Observations from Uganda, where populations located west of the Rift show reduced connectivity with those from eastern regions, support this hypothesis, and similar patterns have been reported in *An. gambiae* (Weedall et al., 2020b). In Kenya, the Rift Valley separates inland populations from coastal regions such as Kilifi, and this large-scale geographic divide likely contributes to restricting gene flow within East Africa (Polo et al., 2025). More broadly, this barrier may underlie the observed differentiation between East African populations and those from West and Central Africa by limiting connectivity across the continent.

Interestingly, summary statistics for genetic diversity also show a difference between the eastern population compared to the others. The Kenyan population exhibited positive Tajima’s D values together with reduced nucleotide diversity and extended runs of homozygosity on chromosome 2RL, consistent with reduced genetic variation and a distinct demographic history. These patterns are typically associated with population bottlenecks, isolation, or recent expansion following a reduction in effective population size. Previous work analysing the same Kenyan population (Polo et al., 2025) has reported similar signals and suggested that they may reflect recent positive selection or demographic changes linked to vector control interventions. These observations highlight that both demographic history and local selective pressures can shape population-specific genetic patterns, and that such regional differences should be considered when interpreting adaptive responses and designing vector control strategies.

Within West Africa, an additional population structure was observed. The population structure analysis all indicated that while some individuals from Saint-Louis and Louga clustered with the Guinea and Nigeria populations, others, particularly from Kolda and Kedougou formed more distinct groups. This pattern may reflect differences in ecological and geographic contexts among the sampling locations. Senegal has several contrasting ecological zones, ranging from Sahelian environments in the north to more humid regions in the south (Alvarez & Govind, 2025; Diouf et al., 2025), which can influence mosquito population structure and local adaptation. In contrast, the Nigerian and Guinean sampling locations share humid environments with perennial river systems that support suitable breeding habitats for *An. funestus*. In Guinea, most mosquitoes were collected in Nzérékoré in the forested southern region of the country, while the Nigerian samples originated from Ogun and Oyo states in southwestern Nigeria, both areas characterized by substantial rainfall and agricultural activity (Adeleke & Olabode, 2025). Such environmental similarities may promote comparable selective pressures and facilitate connectivity between populations. The closer genetic relationship between Guinea and Nigeria may therefore reflect shared ecological conditions and potential ancestral connectivity across the forest and humid savannah belt of West Africa. This highlights the importance of considering regional ecological contexts when interpreting patterns of genetic structure in malaria vectors, as environmental conditions and landscape features can strongly influence mosquito populations.

Consistent with our expectation that strong insecticide pressure leaves detectable genomic footprints, we examined genome-wide signatures of selection and prioritized candidate SNPs potentially driving the selective sweeps detected. The two strongest signals of selection identified in this study were located on chromosome arms 2RL and 3RL, highlighting distinct but complementary adaptive processes shaping *An. funestus* populations. The 2RL region (2RL:8,000,000–9,852,553), encompassing a cluster of detoxification genes, showed the most consistent and pronounced signature of selection across populations. In particular, variants prioritized by iSAFE were shared across all populations and approached fixation in Guinea, Nigeria, and Cameroon, while remaining at lower frequencies in Senegal and Kenya. This pattern is characteristic of an older selective sweep that has spread across multiple populations but is modulated by local ecological conditions or differences in selection intensity (Chotai et al., 2024; Ketchum et al., 2026; Weedall et al., 2020c).

Within this region, the cytochrome P450 gene CYP6A14 emerged as a key candidate. Several high-frequency non-synonymous mutations (L261S, F253I, I94L) were embedded within extended diplotypes showing reduced heterozygosity, suggesting that they are part of dominant selective backgrounds rather than neutral variation. In mosquitoes, cytochrome P450 enzymes are central to metabolic resistance, acting by detoxifying insecticides before they reach their molecular targets, and multiple CYP6 family members have been implicated in resistance in malaria vectors (Riveron et al., 2017, 2018). Although CYP6A14 has been less extensively studied than other P450 genes, evidence from other insect systems suggests that it contributes to xenobiotic detoxification and stress responses (Bacot et al., 2026; Wang et al., 2026; Zhang et al., 2023), raising the possibility that structural changes in this enzyme may enhance its catalytic efficiency. The concordance between iSAFE prioritization, haplotype structure, and allele frequency patterns therefore strongly supports CYP6A14 as a candidate driver of the selective sweep observed in this region. This locus could thus represent an important candidate for further functional investigation to determine whether these mutations modify enzyme activity and contribute to resistance phenotypes.

In contrast, variants identified in the esterase E4-like and sodium/calcium exchanger 3 genes were predicted to have low or modifier effects, yet they co-occurred within the same sweep region (2RL:8,000,000–9,852,553), and were associated with similar haplotypic backgrounds. This suggests that these variants may contribute indirectly to the selective signal, either through regulatory effects or linkage with causal mutations. Importantly, copy number variation was also detected across several detoxification genes within this region, including in LOC125764712 (cytochrome P450 6a8-like), LOC125764714 (probable cytochrome P450 6a14), LOC125764726 (probable cytochrome P450 6a13) and esterases. Gene amplification is a well-established mechanism of metabolic resistance, as increased CNV can elevate enzyme expression and enhance detoxification capacity (Tekoh et al., 2025; Vontas et al., 2020; Wondji et al., 2009). The co-occurrence of SNPs and CNVs within the same genomic region therefore points to a multilayered adaptive architecture, where both sequence variation and structural changes may contribute to the phenotype/s under selection.

Beyond classical detoxification pathways, population-specific signals in this 2RL(2RL:8,000,000– 9,852,553) region point toward additional adaptive mechanisms. In Guinea and Nigeria, variants were detected in loci related to octopamine receptors, including a non-synonymous mutation (G111R) in the β2 receptor that was restricted to Nigerian individuals exhibiting reduced heterozygosity. Octopamine is a key neuromodulator in insects, regulating processes such as locomotion, sensory perception, metabolism, and stress responses (Farooqui, 2012; Roeder, 1999). Because these traits directly influence mosquito survival, host-seeking, and behavior, variation in octopaminergic signalling may represent an alternative route of adaptation distinct from classical metabolic resistance mechanisms. Recent studies have highlighted the importance of octopaminergic signalling in regulating mosquito flight, sensory processing, and host-seeking behaviour. A review of experimental work in *Anopheles* mosquitoes has shown that octopamine modulates the auditory system used during mating swarms, altering the mechanical properties of the mosquito ear and enabling males to detect female flight tones (Freeman et al., 2024; Georgiades et al., 2023). Because successful mating relies on this acoustic communication, disruption of octopaminergic signalling can impair reproductive behaviour and reduce mosquito fitness. More recent studies have further demonstrated that specific octopamine receptors, such as the β2 receptor, regulate mosquito reproductive processes and represent promising targets for population control strategies (Ellis, Bagi, et al., 2025). The presence of selective signals in these genes therefore suggests that, in addition to metabolic detoxification, behavioural and physiological pathways may contribute to how mosquito populations adapt to environmental pressures. Importantly, these pathways are unlikely to confer insecticide resistance directly, but may instead influence traits such as mating success, host interaction, and survival. As such, they represent potential targets for complementary vector control strategies aimed at reducing mosquito fitness or disrupting key behaviours, rather than solely overcoming resistance mechanisms. Understanding the functional role of these pathways could therefore open new avenues for the development of next-generation interventions targeting neural signalling systems in malaria vectors.

A second major signal of selection was identified on chromosome arm 3RL (3RL:12,761,557– 14,664,404), centred on the GABA receptor region. In contrast to the broad and shared signal observed on 2RL, the 3RL sweep appeared more localized, with stronger signals in Nigeria and Cameroon. Diplotype clustering revealed reduced heterozygosity in a subset of Nigerian individuals, within which a non-synonymous mutation (T345S) was detected in the GABA receptor. Although this variant did not reach fixation and showed moderate iSAFE support, its presence within low-diversity haplotypes suggests that it may contribute to the selective signal in this population. The GABA receptor is a well-established target of neurotoxic insecticides (Al Naggar et al., 2025; Gholizadeh et al., 2010), and mutations in this pathway are known to confer resistance by altering receptor sensitivity (Nagi et al., 2025; Wondji et al., 2011). While the variant identified here is distinct from classical resistance mutations, its location within this gene raises the possibility that selection may also act on neural signalling components. Interestingly, the concurrent detection of signals in both GABA and octopamine-related genes suggests that neuromodulatory systems may play a broader role in mosquito adaptation than previously appreciated. These pathways are functionally interconnected and regulate neural excitability, behaviour, and physiological responses to stress, indicating that selection may act not only on detoxification mechanisms but also on how mosquitoes perceive and respond to their environment.

## Conclusion

Overall, this study underscores the importance of considering geographic structure and ecological context when evaluating the evolutionary dynamics of *Anopheles funestus* across Africa. The clear genetic differentiation observed between East, Central, and West African populations, alongside finer-scale structure within regions, highlights that population connectivity and local environmental conditions jointly shape adaptive variation. These patterns demonstrate that geographic proximity alone does not predict evolutionary responses, and that ecological factors play a central role in structuring mosquito populations and their adaptation to vector control pressures.

Our findings further demonstrate the heterogeneous nature of adaptive evolution in *An. funestus* under insecticide and environmental pressures. Strong selective signals in detoxification pathways, particularly within the 2RL cytochrome P450 cluster, were shared across multiple populations, indicating the spread of adaptive alleles under sustained selection. In contrast, localized signals in neuromodulatory loci, including the GABA receptor and octopamine-related genes, highlight population-specific adaptive responses and reveal that additional biological systems beyond metabolic resistance contribute to mosquito adaptation.

These results emphasize the need for region-specific vector control strategies that account for both shared and local patterns of adaptation. Effective resistance management will require continued genomic surveillance, integration of ecological and behavioural data, and expansion beyond classical resistance markers to include alternative pathways influencing mosquito fitness and behaviour. Importantly, the novel signals identified here, spanning structural variants, neuromodulatory loci, and population-specific sweep regions, would not have been detectable through targeted sequencing approaches, underscoring the necessity of whole-genome sequencing for comprehensive surveillance and for broadening our understanding of adaptation in malaria vectors. Such approaches will be critical for developing targeted, sustainable, and next-generation vector control interventions.

## Acknowledgements

This study was supported by the MalariaGEN Vector Observatory which is an international collaboration working to build capacity for malaria vector genomic research and surveillance, and involves contributions by the following institutions and teams. Liverpool School of Tropical Medicine: Kelly Bennett, Victoria Simpson, Lee Hart, Eric Lucas, Martin Donnelly; Wellcome Sanger Institute, Alistair Miles, Eleanor Drury, Mara Lawniczak; The authors would like to thank the staff of the Wellcome Sanger Institute Sample Logistics, Sequencing and Informatics facilities for their contributions. The MalariaGEN Vector Observatory is supported by multiple institutes and funders. The Liverpool School of Tropical Medicine’s participation was supported by the Gates Foundation (INV-068808), by the National Institute of Allergy and Infectious Diseases ([NIAID] R01-AI116811), and additional support from the Medical Research Council (MR/P02520X/1). The latter grant is a UK-funded award and is part of the EDCTP2 programme supported by the European Union. Martin Donnelly is supported by a Royal Society Wolfson Fellowship (RSWF\FT\180003).

At Medical Research Council Unit The Gambia at LSHTM, this study was supported by the Pan-African Malaria Genetic Epidemiology Network (PAMGEN) Human Hereditary and Health is Africa award (H3A/15/002) from African Academy of Science/Science for Africa program, Genomic Surveillance of Malaria in West Africa (GSM, NIHR134717) award from National Institute of Health Research (NIHR), and a BMGF PAMCA grant.

## Data Accessibility and Benefit-Sharing

This study utilises publicly available whole-genome sequence data from the MalariaGEN Anopheles funestus genomic surveillance project, accessible through the malariagen_data API (https://malariagen.github.io/malariagen-data-python/latest/Af1.html). Data were obtained and used in accordance with the MalariaGEN data sharing policies and terms of use.

## Code availability

All custom scripts and analysis code used in this study are available on GitHub at: https://github.com/dany-gaga/an_funestus_genomic_signatures.git

## Author Contributions

H.M.S, B.S.A and A.A.N conceived of and designed the study. H.M.S analysed and interpreted the sequencing data, with input from A.S, M.A, A.A.N and B.S.A. K.L.B and C.S.C reviewed the analysis and contributed to its interpretation. J.B, A.H.K, K.L.B, and C.S.C coordinated data production and curation. O.O, J.M, E.K.L, C.A.N, E.H.A.N, E.O, A.E and E.K.L contributed to sample and data collection. H.M.S. wrote the initial manuscript. H.M.S., A.A.N., B.S.A., K.L.B. and C.S.C. revised the manuscript. All authors provided critical revision of the paper and had full access to all data in the study.

## Supplementary files

**Supplementary file 1:** Summary of sample collection by location, date, and number of individual

**Supplementary file 2**: Population structure within West African *An. funestus* populations Principal component analysis (PCA) based on variants from the inversion-free region of chromosome arm 2RL (57,604,655–90,000,000), showing the first two principal components. Samples are colored by collection site within West Africa (Senegal, Guinea, and Nigeria).

**Supplementary file 3:** Genome-wide selection scans (H12) across chromosome arms for individual sampling sites

Genome-wide selection scans (GWSS) based on the H12 statistic for *An. funestus* populations at individual sampling sites with at least 15 samples. Each panel represents a different geographic location, and plots show window-based H12 values across chromosome arms 2RL (A) 3RL (B) and X (C). Peaks indicate regions of elevated haplotype homozygosity consistent with recent positive selection. Gene annotations are displayed alongside each chromosome to highlight the genomic context of candidate regions.

**Supplementary file 4**: Genes and annotations in candidate regions under selection

**Supplementary file 5:** Patterns of genetic differentiation across candidate genomic regions.

Genomic windows showing pairwise population differentiation (FST) and shared haplotype structure (H1X) across selected regions identified by genome-wide scans. For each region, the upper panel displays FST values along genomic coordinates, with the underlying gene annotation track indicating the position of candidate genes. The corresponding heatmap summarizes pairwise H1X values among populations (Senegal, Guinea, Nigeria, Cameroon, and Kenya), where higher values indicate greater sharing of the most frequent haplotype between population pairs, consistent with a common selective sweep.

**Supplementary file 6**: Allele frequencies of high iSAFE-scoring SNPs identified across all populations

**Supplementary file 7**: Allele frequencies of high iSAFE-scoring SNPs identified within populations

***Supplementary file* 8:** Diplotype clustering within the cytochrome P450 gene cluster on chromosome arm 2RL

Hierarchical clustering of diplotypes across the candidate sweep region chromosome arm 2RL (2RL:8591077-8713423) containing the cytochrome P450 gene cluster The dendrograms represent genetic distances among haplotypes, with individuals colored according to country of origin (Senegal, Cameroon, Nigeria, Guinea, and Kenya). The heterozygosity track below the dendrogram shows patterns of variation across individuals. Additional tracks indicate CNVs per genes.

**Supplementary file 9:** Diplotype clustering within the octopamine receptor and GABA receptor loci on chromosome arms 2RL and 3RL

Hierarchical clustering of diplotypes across the candidate sweep region on (A) chromosome arm 2RL (2RL:8915154-9064309) containing the octopamine genes and (B) chromosome arm 3RL (3 RL:13516317-13612792) containing the GABA receptor locus. The dendrograms represent genetic distances among haplotypes, with individuals colored according to country of origin (Senegal, Cameroon, Nigeria, Guinea, and Kenya). The heterozygosity track below the dendrogram shows patterns of variation across individuals. Additional tracks indicate CNVs per genes.

