## Supplementary file 1 for "Genomic signatures of selection in *Anopheles funestus* reveal shared and population-specific adaptive variation across African populations"

**Supplementary file 1:** Summary of sample collection by location, date, and number of individual

| Year | Month | Country | Sites of collection | Latitude | Longitude | Number of samples |
|---|---|---|---|---|---|---|
| *2020* | 7 | Senegal | Kaolack | 13.784 | -16.19 | 1 |
| *2020* | 7 | Senegal | Kedougou | 12.553 | -12.529 | 4 |
| *2020* | 7 | Senegal | Kedougou | 12.623 | -11.61 | 1 |
| *2020* | 7 | Senegal | Kolda | 12.832 | -15.005 | 5 |
| *2021* | 9 | Senegal | Kolda | 12.832 | -15.005 | 16 |
| *2022* | 10 | Senegal | Kolda | 12.832 | -15.005 | 19 |
| *2022* | 10 | Senegal | Louga | 15.953 | -15.921 | 13 |
| *2022* | 10 | Senegal | Saint Louis | 16.325 | -15.767 | 1 |
| *2022* | 10 | Senegal | Saint Louis | 16.542 | -14.801 | 11 |
| *2020* | 9 | Cameroon | South | 2.821 | 10.135 | 81 |
| *2018* | 8 | Nigeria | Ogun | 7.361 | 3.6 | 18 |
| *2018* | 8 | Nigeria | Ogun | 7.42 | 3.64 | 84 |
| *2018* | 9 | Nigeria | Ogun | 7.235 | 3.749 | 20 |
| *2018* | 9 | Nigeria | Ogun | 7.361 | 3.6 | 87 |
| *2018* | 9 | Nigeria | Ogun | 7.42 | 3.64 | 27 |
| *2018* | 9 | Nigeria | Oyo | 7.445 | 3.633 | 27 |
| *2022* | 1 | Guinea | Kankan | 11.37 | -9.15 | 7 |
| *2022* | 1 | Guinea | Kankan | 11.416 | -9.058 | 6 |
| *2022* | 1 | Guinea | Nzerekore | 7.846 | -8.846 | 88 |
| *2022* | 1 | Guinea | Nzerekore | 8.591 | -9.959 | 95 |
| *2010* | -1 | Kenya | Kilifi | -3.511 | 39.909 | 2 |
| *2012* | -1 | Kenya | Kilifi | -3.511 | 39.909 | 22 |
