## Supplementary file 2 for "Genomic signatures of selection in *Anopheles funestus* reveal shared and population-specific adaptive variation across African populations"

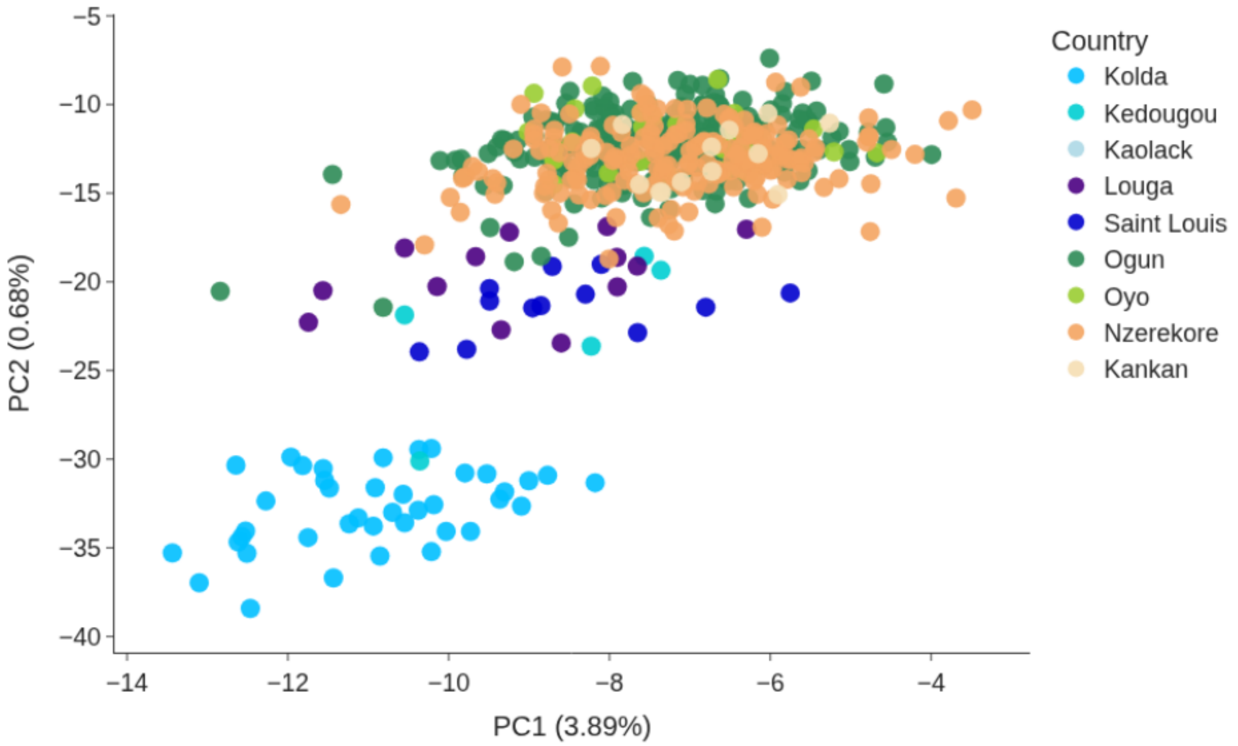


**Supplementary file 2**: Population structure within West African *An. funestus* populations Principal component analysis (PCA) based on variants from the inversion-free region of chromosome arm 2RL (57,604,655–90,000,000), showing the first two principal components. Samples are colored by collection site within West Africa (Senegal, Guinea, and Nigeria).
