## Supplementary file 3 for "Genomic signatures of selection in *Anopheles funestus* reveal shared and population-specific adaptive variation across African populations"

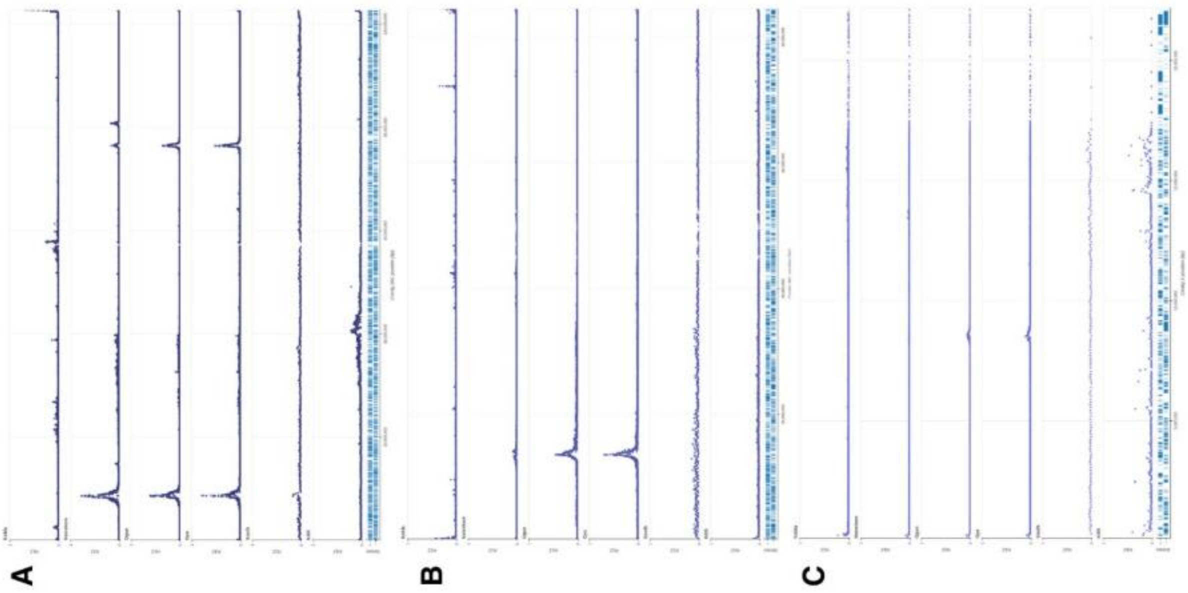

**Supplementary file 3:** Genome-wide selection scans (H12) across chromosome arms for individual sampling sites

Genome-wide selection scans (GWSS) based on the H12 statistic for *An. funestus* populations at individual sampling sites with at least 15 samples. Each panel represents a different geographic location, and plots show window-based H12 values across chromosome arms 2RL (A) 3RL (B) and X (C). Peaks indicate regions of elevated haplotype homozygosity consistent with recent positive selection. Gene annotations are displayed alongside each chromosome to highlight the genomic context of candidate regions.
