## Supplementary file 5 for "Genomic signatures of selection in *Anopheles funestus* reveal shared and population-specific adaptive variation across African populations"

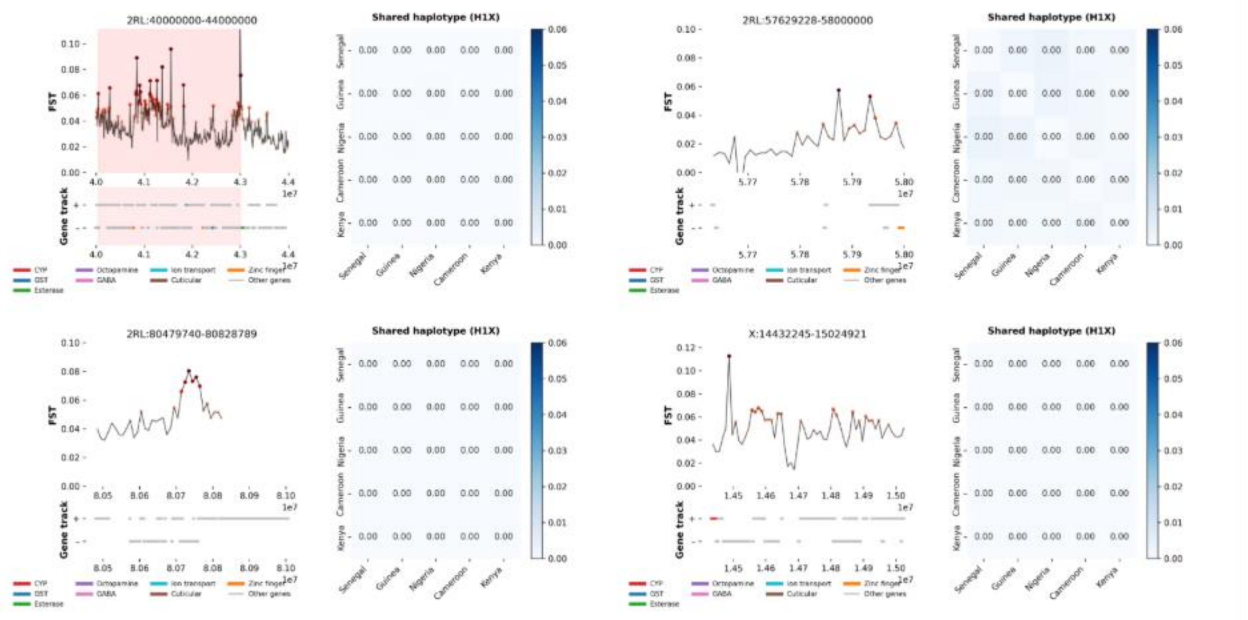


**Supplementary file 5:** Patterns of genetic differentiation across candidate genomic regions.

Genomic windows showing pairwise population differentiation (FST) and shared haplotype structure (H1X) across selected regions identified by genome-wide scans. For each region, the upper panel displays FST values along genomic coordinates, with the underlying gene annotation track indicating the position of candidate genes. The corresponding heatmap summarizes pairwise H1X values among populations (Senegal, Guinea, Nigeria, Cameroon, and Kenya), where higher values indicate greater sharing of the most frequent haplotype between population pairs, consistent with a common selective sweep.
