## Supplementary file 8 for "Genomic signatures of selection in *Anopheles funestus* reveal shared and population-specific adaptive variation across African populations"

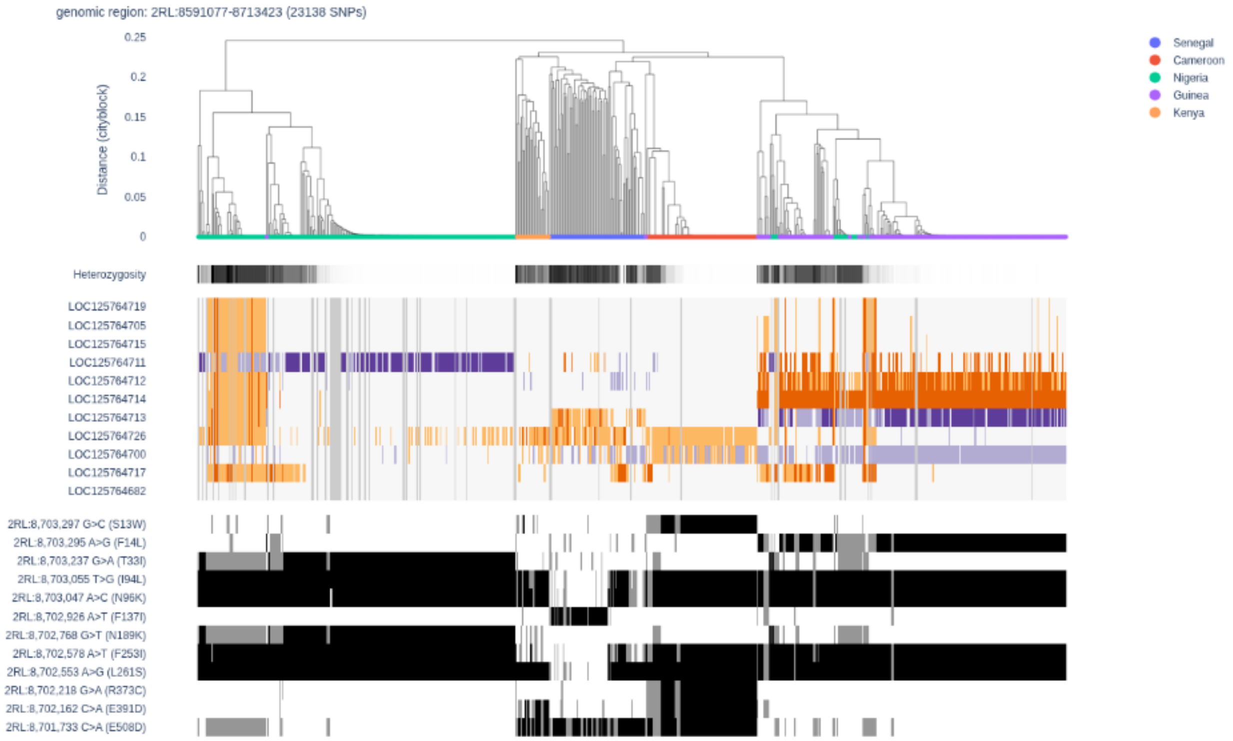


**Supplementary file 8:** Diplotype clustering within the cytochrome P450 gene cluster on chromosome arm 2RL

Hierarchical clustering of diplotypes across the candidate sweep region chromosome arm 2RL (2RL:8591077-8713423) containing the cytochrome P450 gene cluster The dendrograms represent genetic distances among haplotypes, with individuals colored according to country of origin (Senegal, Cameroon, Nigeria, Guinea, and Kenya). The heterozygosity track below the dendrogram shows patterns of variation across individuals. Additional tracks indicate CNVs per genes.
