## Supplementary file 9 for "Genomic signatures of selection in *Anopheles funestus* reveal shared and population-specific adaptive variation across African populations"

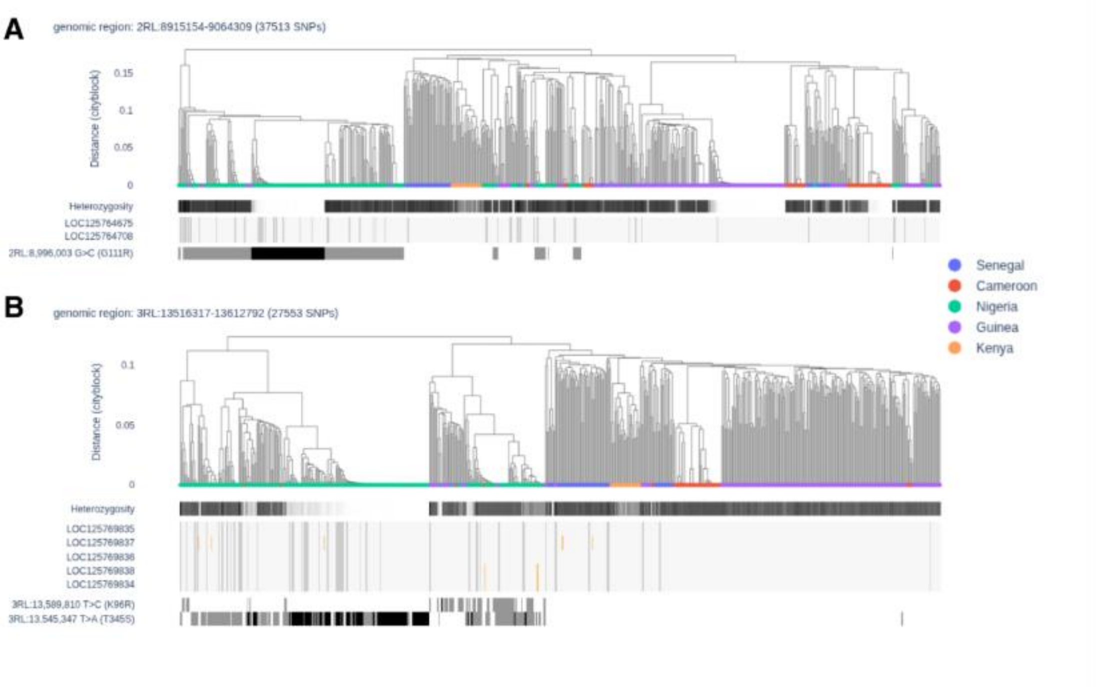


**Supplementary file 9:** Diplotype clustering within the octopamine receptor and GABA receptor loci on chromosome arms 2RL and 3RL

Hierarchical clustering of diplotypes across the candidate sweep region on (A) chromosome arm 2RL (2RL:8915154-9064309) containing the octopamine genes and (B) chromosome arm 3RL (3 RL:13516317-13612792) containing the GABA receptor locus. The dendrograms represent genetic distances among haplotypes, with individuals colored according to country of origin (Senegal, Cameroon, Nigeria, Guinea, and Kenya). The heterozygosity track below the dendrogram shows patterns of variation across individuals. Additional tracks indicate CNVs per genes.
