## Supplementary file 6 for "Genomic signatures of selection in *Anopheles funestus* reveal shared and population-specific adaptive variation across African populations"

| Region | Transcript | Genomic position | iSAFE (All countries) | SNP frequency in Senegal | SNP frequency in Guinea | SNP frequency in Nigeria | SNP frequency in Cameroon | SNP frequency in Kenya | Variant effect | Variant annotation | Description |
|---|---|---|---|---|---|---|---|---|---|---|---|
| 2RL:8000000-9852553 | LOC125764705_t1 | 8703059 | 0.748 | 0.29 | 0.98 | 1.00 | 0.98 | 0.88 | SYNONYMOUS_CODING | 2RL:8,703,059 A>C (R92R) | probable cytochrome P450 6a14 |
| 2RL:8000000-9852553 | LOC125764682_t3 | 8703059 | 0.748 | 0.29 | 0.98 | 1.00 | 0.98 | 0.88 | TRANSCRIPT | 2RL:8,703,059 A>C | sodium/calcium exchanger 3, transcript variant X7 |
| 2RL:8000000-9852553 | LOC125764682_t3 | 8689600 | 0.738 | 0.00 | 0.00 | 0.00 | 0.00 | 0.00 | TRANSCRIPT | 2RL:8,689,600 T>A | sodium/calcium exchanger 3, transcript variant X7 |
| 2RL:8000000-9852553 | LOC125764713_t1 | 8689600 | 0.738 | 0.00 | 0.00 | 0.00 | 0.00 | 0.00 | INTRONIC | 2RL:8,689,600 T>A | probable cytochrome P450 6a14, transcript variant X2 |
| 2RL:8000000-9852553 | LOC125764705_t1 | 8703047 | 0.726 | 0.32 | 0.98 | 1.00 | 0.98 | 0.94 | NON_SYNONYMOUS_CODING | 2RL:8,703,047 A>C (N96K) | probable cytochrome P450 6a14 |
| 2RL:8000000-9852553 | LOC125764682_t3 | 8703047 | 0.726 | 0.32 | 0.98 | 1.00 | 0.98 | 0.94 | TRANSCRIPT | 2RL:8,703,047 A>C | sodium/calcium exchanger 3, transcript variant X7 |
| 2RL:8000000-9852553 | LOC125764682_t3 | 8622756 | 0.522 | 0.39 | 0.92 | 0.92 | 0.99 | 0.12 | TRANSCRIPT | 2RL:8,622,756 A>C | sodium/calcium exchanger 3, transcript variant X7 |
| 2RL:8000000-9852553 | LOC125764682_t3 | 8709861 | 0.510 | 0.52 | 0.98 | 0.98 | 1.00 | 0.46 | TRANSCRIPT | 2RL:8,709,861 A>C | sodium/calcium exchanger 3, transcript variant X7 |
| 2RL:8000000-9852553 | LOC125764682_t3 | 8591332 | 0.503 | 0.01 | 0.83 | 0.89 | 0.06 | 0.00 | FIVE_PRIME_UTR | 2RL:8,591,332 T>A | sodium/calcium exchanger 3, transcript variant X7 |
| 2RL:8000000-9852553 | LOC125764734_t1 | 8718454 | 0.492 | 0.02 | 0.84 | 0.91 | 0.04 | 0.00 | INTRONIC | 2RL:8,718,454 C>A | integumentary mucin A.1-like |
| 2RL:8000000-9852553 | LOC125764682_t3 | 8653065 | 0.484 | 0.68 | 0.99 | 0.99 | 0.98 | 0.31 | TRANSCRIPT | 2RL:8,653,065 G>A | sodium/calcium exchanger 3, transcript variant X7 |
| 2RL:8000000-9852553 | LOC125764682_t3 | 8710475 | 0.480 | 0.00 | 0.85 | 0.86 | 0.04 | 0.00 | THREE_PRIME_UTR | 2RL:8,710,475 T>A | sodium/calcium exchanger 3, transcript variant X7 |
| 2RL:8000000-9852553 | LOC125764682_t3 | 8635968 | 0.480 | 0.55 | 0.96 | 0.98 | 0.99 | 0.44 | TRANSCRIPT | 2RL:8,635,968 G>A | sodium/calcium exchanger 3, transcript variant X7 |
| 2RL:8000000-9852553 | LOC125764682_t3 | 8669329 | 0.472 | 0.06 | 0.90 | 0.92 | 0.04 | 0.08 | TRANSCRIPT | 2RL:8,669,329 G>A | sodium/calcium exchanger 3, transcript variant X7 |
| 2RL:8000000-9852553 | LOC125761417_t1 | 8537263 | 0.462 | 0.04 | 0.80 | 0.85 | 0.05 | 0.00 | TRANSCRIPT | 2RL:8,537,263 C>A | sodium/potassium-transporting ATPase subunit alpha, transcript variant X11 |
| 3RL:12761557-14664404 | LOC125769835_t1 | 13603457 | 0.350 | 0.00 | 0.12 | 0.84 | 0.46 | 0.00 | TRANSCRIPT | 3RL:13,603,457 G>A | gamma-aminobutyric acid receptor subunit beta, transcript variant X11 |
| 3RL:12761557-14664404 | LOC125769834_t1 | 13519933 | 0.317 | 0.01 | 0.00 | 0.00 | 0.00 | 0.00 | INTRONIC | 3RL:13,519,933 G>A | probable nucleoporin Nup54 |
| 3RL:12761557-14664404 | LOC125769838_t1 | 13519933 | 0.317 | 0.01 | 0.00 | 0.00 | 0.00 | 0.00 | SPLICE_REGION | 3RL:13,519,933 G>A | uncharacterized LOC125769838 |
| 3RL:12761557-14664404 | LOC125769835_t1 | 13547718 | 0.309 | 0.00 | 0.01 | 0.70 | 0.35 | 0.00 | TRANSCRIPT | 3RL:13,547,718 G>A | gamma-aminobutyric acid receptor subunit beta, transcript variant X11 |
| 3RL:12761557-14664404 | LOC125769834_t1 | 13516501 | 0.307 | 0.01 | 0.11 | 0.76 | 0.48 | 0.08 | SYNONYMOUS_CODING | 3RL:13,516,501 G>A (D602D) | probable nucleoporin Nup54 |
| 3RL:12761557-14664404 | LOC125769835_t1 | 13547968 | 0.306 | 0.00 | 0.02 | 0.70 | 0.40 | 0.00 | TRANSCRIPT | 3RL:13,547,968 G>A | gamma-aminobutyric acid receptor subunit beta, transcript variant X11 |
| 3RL:12761557-14664404 | LOC125769835_t1 | 13552755 | 0.305 | 0.00 | 0.01 | 0.68 | 0.36 | 0.00 | TRANSCRIPT | 3RL:13,552,755 A>C | gamma-aminobutyric acid receptor subunit beta, transcript variant X11 |
| 3RL:12761557-14664404 | LOC125769835_t1 | 13557381 | 0.305 | 0.00 | 0.01 | 0.68 | 0.36 | 0.00 | TRANSCRIPT | 3RL:13,557,381 A>C | gamma-aminobutyric acid receptor subunit beta, transcript variant X11 |
| 3RL:12761557-14664404 | LOC125769835_t1 | 13552606 | 0.301 | 0.00 | 0.02 | 0.68 | 0.36 | 0.00 | TRANSCRIPT | 3RL:13,552,606 A>C | gamma-aminobutyric acid receptor subunit beta, transcript variant X11 |
| 3RL:12761557-14664404 | LOC125769835_t1 | 13554611 | 0.296 | 0.01 | 0.02 | 0.68 | 0.36 | 0.00 | TRANSCRIPT | 3RL:13,554,611 G>A | gamma-aminobutyric acid receptor subunit beta, transcript variant X11 |
| 3RL:12761557-14664404 | LOC125772323_t1 | 13503573 | 0.296 | 0.00 | 0.04 | 0.63 | 0.43 | 0.00 | INTRONIC | 3RL:13,503,573 G>A | uncharacterized LOC125772323 |
| 3RL:12761557-14664404 | LOC125769835_t1 | 13569844 | 0.296 | 0.00 | 0.01 | 0.67 | 0.33 | 0.00 | TRANSCRIPT | 3RL:13,569,844 T>A | gamma-aminobutyric acid receptor subunit beta, transcript variant X11 |
| 3RL:12761557-14664404 | LOC125769835_t1 | 13563506 | 0.296 | 0.00 | 0.02 | 0.67 | 0.34 | 0.00 | TRANSCRIPT | 3RL:13,563,506 A>C | gamma-aminobutyric acid receptor subunit beta, transcript variant X11 |
| 3RL:12761557-14664404 | LOC125769835_t1 | 13571797 | 0.296 | 0.00 | 0.01 | 0.67 | 0.33 | 0.00 | TRANSCRIPT | 3RL:13,571,797 A>C | gamma-aminobutyric acid receptor subunit beta, transcript variant X11 |
| 3RL:12761557-14664404 | LOC125769835_t1 | 13574693 | 0.296 | 0.00 | 0.01 | 0.67 | 0.33 | 0.00 | TRANSCRIPT | 3RL:13,574,693 G>A | gamma-aminobutyric acid receptor subunit beta, transcript variant X11 |
