## Supplementary file 7 for "Genomic signatures of selection in *Anopheles funestus* reveal shared and population-specific adaptive variation across African populations"

| Region | Transcript | Genomic position | iSAFE (max) | SNP frequency in Senegal | SNP frequency in Guinea | SNP frequency in Nigeria | SNP frequency in Cameroon | SNP frequency in Kenya | Variant effect | Variant annotation | Description |
|---|---|---|---|---|---|---|---|---|---|---|---|
| 2RL:8000000-9852553 | LOC125764708_t3 | 8936172 | 0.451 | 0.11 | 0.09 | 0.57 | 0.61 | 0.00 | TRANSCRIPT | 2RL:8,936,172 G>A | octopamine receptor beta-2R, transcript variant X4 |
| 2RL:8000000-9852553 | LOC125764675_t1 | 9014853 | 0.404 | 0.17 | 0.04 | 0.51 | 0.61 | 0.04 | INTRONIC | 2RL:9,014,853 G>A | uncharacterized LOC125764675 |
| 2RL:8000000-9852553 | LOC125764708_t3 | 8977768 | 0.393 | 0.09 | 0.05 | 0.52 | 0.60 | 0.08 | TRANSCRIPT | 2RL:8,977,768 G>A | octopamine receptor beta-2R, transcript variant X4 |
| 2RL:8000000-9852553 | LOC125764708_t3 | 8927915 | 0.366 | 0.07 | 0.55 | 0.55 | 0.02 | 0.00 | TRANSCRIPT | 2RL:8,927,915 G>A | octopamine receptor beta-2R, transcript variant X4 |
| 2RL:8000000-9852553 | LOC125764708_t3 | 8944108 | 0.366 | 0.00 | 0.00 | 0.00 | 0.00 | 0.00 | TRANSCRIPT | 2RL:8,944,108 A>C | octopamine receptor beta-2R, transcript variant X4 |
| 2RL:8000000-9852553 | LOC125764675_t1 | 9058990 | 0.357 | 0.00 | 0.00 | 0.00 | 0.00 | 0.00 | INTRONIC | 2RL:9,058,990 C>A | uncharacterized LOC125764675 |
| 2RL:8000000-9852553 | LOC125764675_t1 | 9014380 | 0.352 | 0.00 | 0.00 | 0.00 | 0.00 | 0.00 | INTRONIC | 2RL:9,014,380 A>C | uncharacterized LOC125764675 |
| 2RL:8000000-9852553 | LOC125764675_t1 | 9011162 | 0.347 | 0.00 | 0.52 | 0.04 | 0.48 | 0.00 | INTRONIC | 2RL:9,011,162 G>A | uncharacterized LOC125764675 |
| 2RL:8000000-9852553 | LOC125764708_t3 | 8939310 | 0.332 | 0.06 | 0.05 | 0.52 | 0.49 | 0.40 | TRANSCRIPT | 2RL:8,939,310 G>A | octopamine receptor beta-2R, transcript variant X4 |
| 2RL:8000000-9852553 | LOC125764708_t3 | 8939312 | 0.332 | 0.08 | 0.05 | 0.51 | 0.49 | 0.40 | TRANSCRIPT | 2RL:8,939,312 G>A | octopamine receptor beta-2R, transcript variant X4 |
| 2RL:76000000-76838987 | LOC125763932_t3 | 76594263 | 0.468 | 0.01 | 0.00 | 0.00 | 0.00 | 0.00 | FIVE_PRIME_UTR | 2RL:76,594,263 G>A | uncharacterized protein DDB_G0271670-like, transcript variant X2 |
| 2RL:76000000-76838987 | LOC125763932_t3 | 76594015 | 0.466 | 0.00 | 0.00 | 0.00 | 0.00 | 0.00 | FIVE_PRIME_UTR | 2RL:76,594,015 G>A | uncharacterized protein DDB_G0271670-like, transcript variant X2 |
| 2RL:76000000-76838987 | LOC125763932_t3 | 76597916 | 0.464 | 0.00 | 0.00 | 0.00 | 0.00 | 0.00 | TRANSCRIPT | 2RL:76,597,916 T>A | uncharacterized protein DDB_G0271670-like, transcript variant X2 |
| 2RL:76000000-76838987 | LOC125763932_t3 | 76595833 | 0.464 | 0.00 | 0.22 | 0.47 | 0.02 | 0.00 | TRANSCRIPT | 2RL:76,595,833 T>A | uncharacterized protein DDB_G0271670-like, transcript variant X2 |
| 2RL:76000000-76838987 | LOC125763932_t3 | 76598356 | 0.464 | 0.00 | 0.00 | 0.00 | 0.00 | 0.00 | TRANSCRIPT | 2RL:76,598,356 G>A | uncharacterized protein DDB_G0271670-like, transcript variant X2 |
| 2RL:76000000-76838987 | LOC125763932_t3 | 76598753 | 0.464 | 0.00 | 0.00 | 0.00 | 0.00 | 0.00 | TRANSCRIPT | 2RL:76,598,753 G>A | uncharacterized protein DDB_G0271670-like, transcript variant X2 |
| 2RL:76000000-76838987 | LOC125763932_t3 | 76597980 | 0.464 | 0.00 | 0.00 | 0.00 | 0.00 | 0.00 | TRANSCRIPT | 2RL:76,597,980 A>C | uncharacterized protein DDB_G0271670-like, transcript variant X2 |
| 2RL:76000000-76838987 | LOC125763932_t3 | 76602753 | 0.461 | 0.00 | 0.00 | 0.00 | 0.00 | 0.00 | TRANSCRIPT | 2RL:76,602,753 C>A | uncharacterized protein DDB_G0271670-like, transcript variant X2 |
| 2RL:76000000-76838987 | LOC125763932_t3 | 76605501 | 0.461 | 0.00 | 0.00 | 0.00 | 0.00 | 0.00 | TRANSCRIPT | 2RL:76,605,501 A>C | uncharacterized protein DDB_G0271670-like, transcript variant X2 |
| 2RL:76000000-76838987 | LOC125763932_t3 | 76599104 | 0.461 | 0.00 | 0.00 | 0.00 | 0.00 | 0.00 | TRANSCRIPT | 2RL:76,599,104 A>C | uncharacterized protein DDB_G0271670-like, transcript variant X2 |
| 2RL:80479740-80828789 | LOC125761952_t3 | 80764400 | 0.325 | 0.02 | 0.38 | 0.04 | 0.20 | 0.00 | TRANSCRIPT | 2RL:80,764,400 C>A | ATP-binding cassette sub-family C member Sur, transcript variant X1 |
| 2RL:80479740-80828789 | LOC125761964_t1 | 80701303 | 0.321 | 0.01 | 0.38 | 0.02 | 0.18 | 0.00 | INTRONIC | 2RL:80,701,303 G>A | carbohydrate sulfotransferase 1 |
| 2RL:80479740-80828789 | LOC125761962_t1 | 80710294 | 0.311 | 0.00 | 0.00 | 0.00 | 0.00 | 0.00 | THREE_PRIME_UTR | 2RL:80,710,294 A>C | probable ATP-dependent RNA helicase DDX47 |
| 2RL:80479740-80828789 | LOC125761956_t1 | 80732869 | 0.294 | 0.00 | 0.00 | 0.00 | 0.00 | 0.00 | TRANSCRIPT | 2RL:80,732,869 A>C | AF4/FMR2 family member lilli, transcript variant X1 |
